# A Whale With A History: Sighting Twain The Humpback Over Three Decades

**DOI:** 10.1101/2022.10.07.511372

**Authors:** James P. Crutchfield, Alexandra M. Jurgens, Janice M. Straley, Ted Cheeseman

**Affiliations:** Complexity Sciences Center, Physics and Astronomy Department, University of California at Davis, Davis, California 95616, USA; College of Fisheries and Ocean Sciences, University of Alaska Southeast, Sitka Campus, Sitka, Alaska 99835, USA; HappyWhale.com, Santa Cruz, California, USA; Marine Ecology Research Centre, Southern Cross University, Lismore Australia

**Keywords:** Megaptera novaeangliae, humpback whale, nonhuman communication, interspecies communication, animal vocalization, mirror study, ocean acoustics

## Abstract

Extended acoustic interactions with a humpback whale (*Megaptera novaeangliae*) were captured via playbacks of the purported “throp” social call and hydrophone recordings of the animal’s vocalized responses during August 2021 in Frederick Sound, Southeast Alaska. Fluke photographs identified the animal as a female named Twain (HappyWhale.com identity SEAK-0401) first observed some 38 years ago. We document Twain’s life history via sightings made over almost four decades. The observational history gives illuminating snapshots of the long history of the individual behind the acoustic interactions.

## I. INTRODUCTION

Recently in Southeast Alaska, a scientific team encountered a humpback whale (*Megaptera novaeangliae*) and engaged the animal in a half-hour long exchange of acoustic calls and responses. Apparently, the animal was motivated to communicate. This, in turn, led to an interest in this particular animal—later identified as a female named Twain (HappyWhale.com identity SEAK-0401) first seen in 1984, over a third of a century ago. Reference [1] reports on details of the encounter, her identification, and quantitative analyses of the hydrophone recordings of the extended encounter.

The following complements that report by recounting Twain’s life history via visual observations made over 38 years, presenting as much as is known (and allowed for public distribution). Though episodic, the observations made in Hawaii and Southeast Alaska illuminate the long history of the animal that engaged in the acoustic interactions. Though decidedly incomplete, the almost four decades of documented observations very likely covers the majority of Twain’s life, giving an indirect view into her individuality.

Figures 1 and 2 show Twain’s migration paths between sightings, illustrating the long (≈ 3000 mile) seasonal (Summer-to-Winter and Winter-to-Summer) trajectories typical of her humpback cohort. Figure 2a shows the same but for sightings around west Maui, Hawaii—the cohort’s winter breeding grounds. And, Figure 2b shows the paths in and around Southeast Alaska, largely centered on Frederick Sound—the cohort’s summer feeding grounds.

**FIG. 1:**
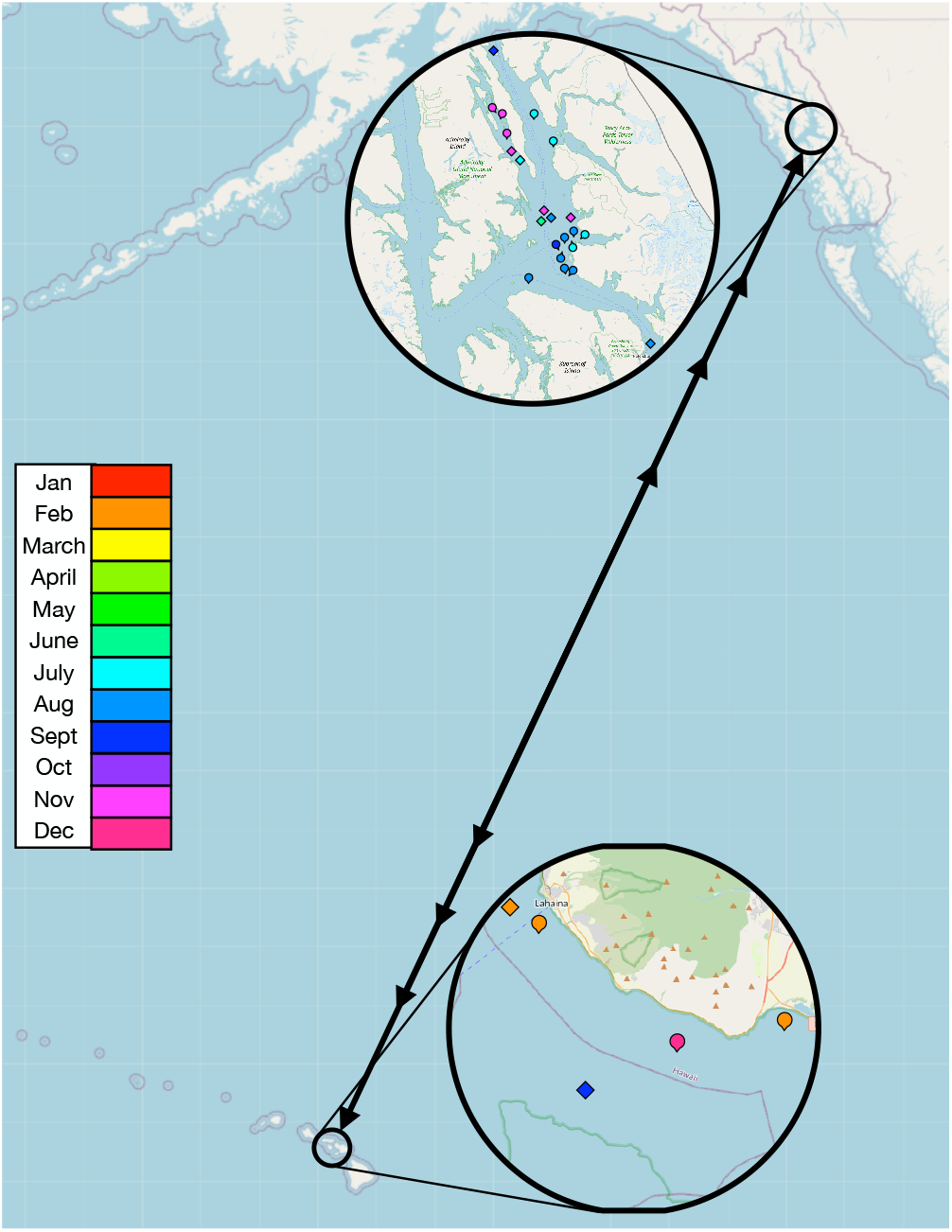
Twain migrations across the north Pacific, seasonally transiting 2, 800 miles between (Summer) southeast Alaska (Fig. 2b) and (Winter) Maui, Hawaii (Fig. 2a). Circles indicate sightings with known latitude-longitude; diamonds those with only region/location. See Table II. OpenStreetMap CC BY-SA 2.0.

**FIG. 2:**
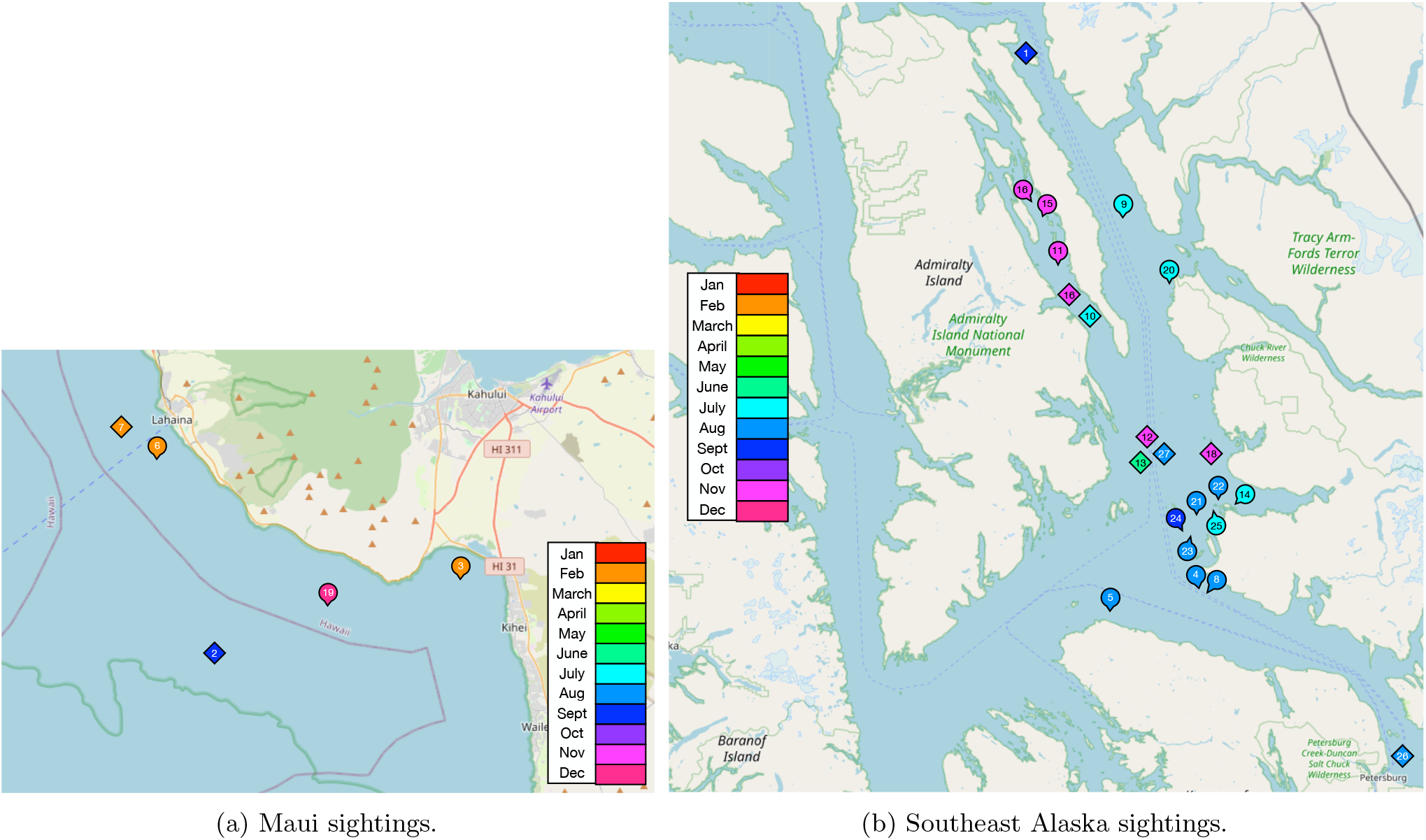
Twain sightings: (a) West Maui and (b) Southeast Alaska. Numbered circles indicate sightings with known latitude-longitude; numbered diamonds those with only region/location, approximately placed. See Table II. OpenStreetMap CC BY-SA 2.0.

Table II lists the over two dozen documented sightings of Twain. Data there include dates, times, locations, and observers. Figures 3 and 4 present a photographic gallery of her flukes—images cataloged and used to identify her over the years. The first set contains historical fluke photos, while the second set documents recent encounters including that of the acoustic interactions, in the last two years.

**FIG. 3:**
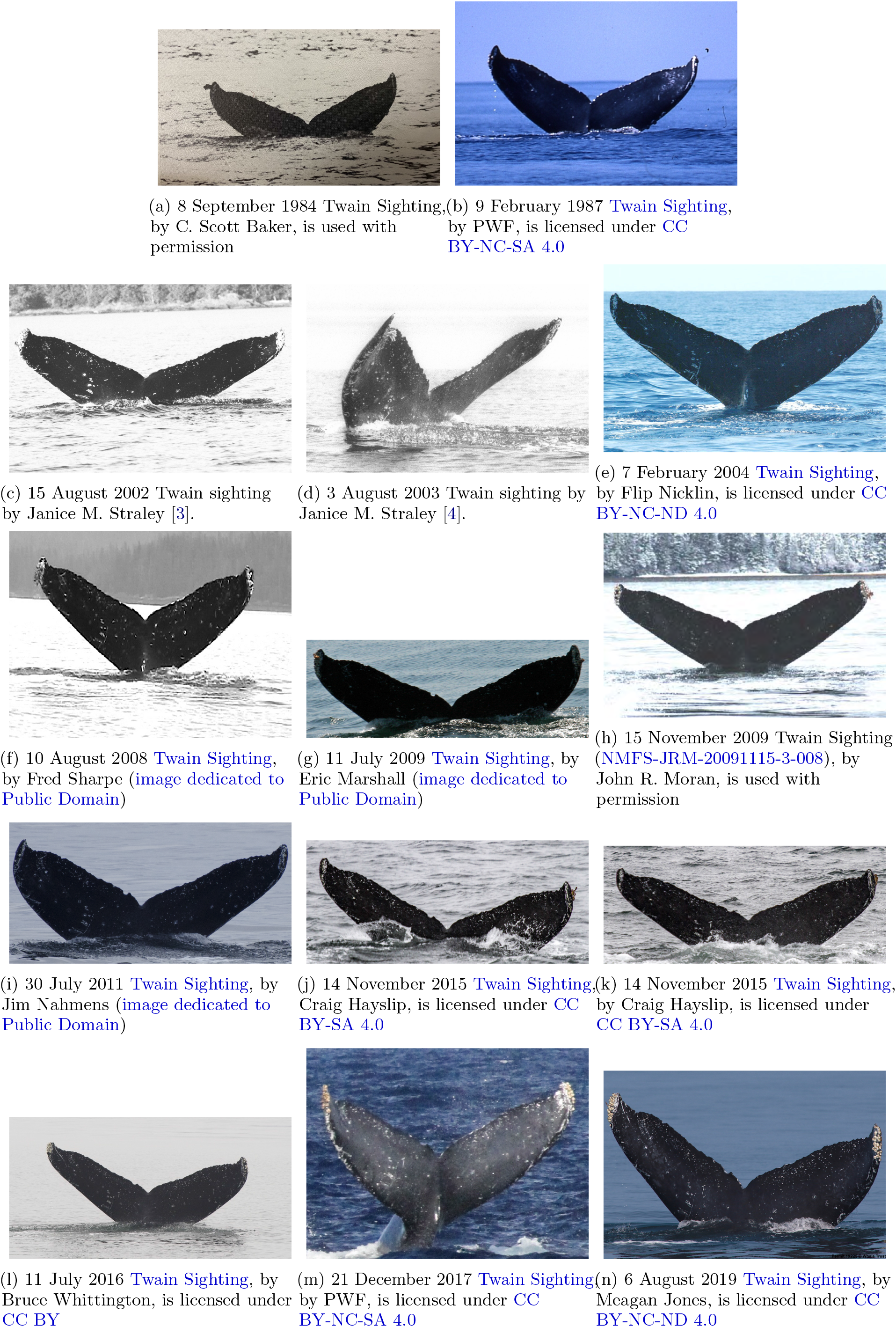
Historical humpback whale Twain (SEAK-0401) identifications in chronological order. First sighting at least as early as 1984 in Alaska; see Fig. 3a. See Table I for alternate identification tags and Table II for sighting details.

**FIG. 4:**
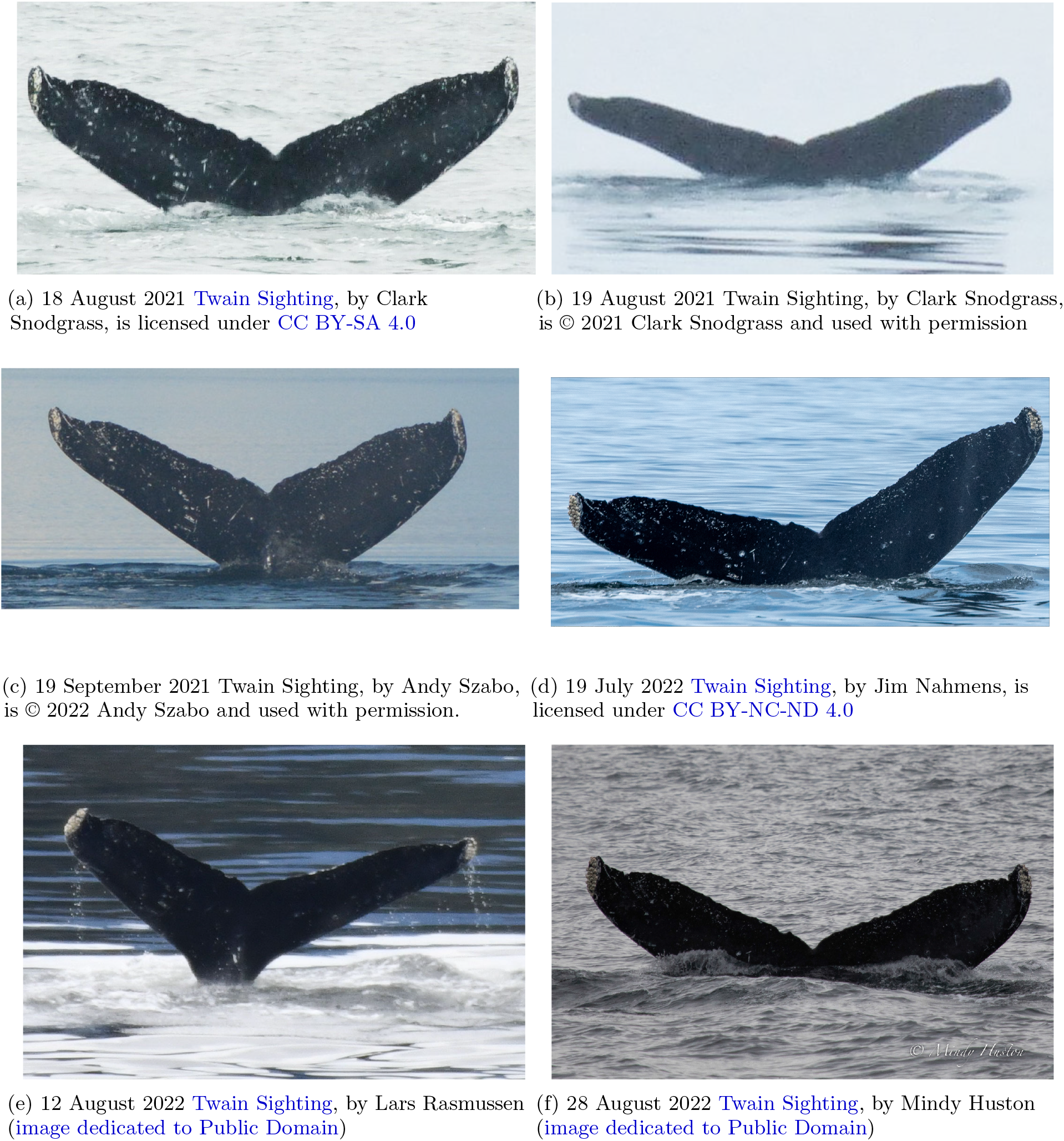
Recent humpback whale Twain (SEAK-0401) sightings (2021 and after): Most recently seen 28 August 2022 in southeast Alaska. See Table I for alternate identification tags and Table II for sighting details. (HappyWhale.com, accessed on or before 16 August 2022.)

The bulk of the data collected here is publicly available and was largely extracted from the online database HappyWhale.com [2] and the UASE 2012 catalog. However, when possible and permitted, it includes observations from other databases and from individual observers.

## II. ACOUSTIC INTERACTIONS

To motivate collecting observations of Twain’s long life history, it will help to summarize the results reported in Ref. [1]. This sets the scientific context and grounds the interest in this individual animal.

Though unidentified at the time, Twain was encountered on 18 August 2021, documented both by fluke photograph (see Fig. 4a) and by hydrophone recording. The most notable event occurred when reviewing the hydrophone recording: there was a singularly clear example of the purported humpback “throp” social call [5–10]. That recording was selected for use in the next day’s acoustic playback experiments.

The extended call-and-response exchanges occurred on 19 August. The exchanges were initiated by the very close approach (20 m) of the animal to the research vessel. Researchers then decided to broadcast the recording of the previous day’s selected call through an underwater loud-speaker. In fact, two additional playbacks were broadcast roughly a minute apart until a response vocalization—a throp social call—was heard from the animal.

Fluke photographs were also taken during the exchanges. Afterwards the animal was identified using HappyWhale.com as SEAK-0404 (Twain) from uploaded fluke photographs. Reference [1]’s statistical analyses of the hydrophone recordings corroborated that the exchanges involved a single individual and that the individual was the animal who’s throp call had been recorded the previous day and used, inadvertently and unintentionally, as the playback call during the exchanges.

Rapid access to the HappyWhale.com online database greatly facilitated identifying the animal encountered. It should also be pointed out that this was also helped in large measure by being in line-of-sight of a land-based cell tower that gave online access to HappyWhale.com. The fluke shots were matched in short order as SEAK-0401, after properly preparing and uploading them.

Reference [1] provides fuller detail. The following complements that report with the long history of Twain’s sightings in Southeast Alaska and Hawaii.

## III. SIGHTINGS

One of the conundrums animal behavior databases such as HappyWhale.com are rapidly coming to address is that a number of observers have seen and photographed the same animal. One consequence is that a given animal becomes known by a large number of identification labels and the supporting observation data is stored across many databases. For example, Table I lists the ten current IDs for Twain. Needless to say, this redundant labeling both aids and complicates confident identification.

**TABLE I:** Diversity of Twain identifying labels. See Table III for abbreviations.

| Organization (Database) | Identification |
| --- | --- |
| UH KBMML | SEAK-0401 |
|  | AK19_0098 |
| UASE/GBNP | FS02-24a-01a |
| UASE/GBNP | FS03-11(34A) |
|  | HI04-0062 |
|  | HW-MN0440644 |
|  | PWF-NP_3005 |
| Oregon State | OSUWTG-MnSEAK-314 |
|  | SPLASH-430588 |
| UASE/GBNP | SEAK-2450 |

Humpback whale morphology cooperates to some extent to ameliorate these problems and improve unique identifications. The ventral side of their flukes carries unique marks—genetically or not determined pigmentations and shapes, cuts, and abrasions due to fishing-gear entanglements or encounters with boats, ships, killer whales, and great white sharks—to mention a few sources of their unique markings. HappyWhale.com takes properly formatted user-uploaded fluke photographs and applies a machine-learning image-classification algorithm to these markings to compare a subject animal to those fluke photos already uploaded and identified. It does so with remarkably high accuracy.

There are other databases than this, though. In many cases, these have been assembled over many years by hand by working marine biologists and citizen scientists alike. One—*Humpback Whales of Southeast Alaska*, that was used here—is maintained by the University of Alaska Southeast (UASE) [11]. And, there are individual whale watchers that curate their own collection of humpback photo IDs.

The result then of combing through these databases and contacting their curators and a number of individual contributors is compiled in Table II. It catalogs most all known sightings of humpback whale Twain.

**TABLE II:**
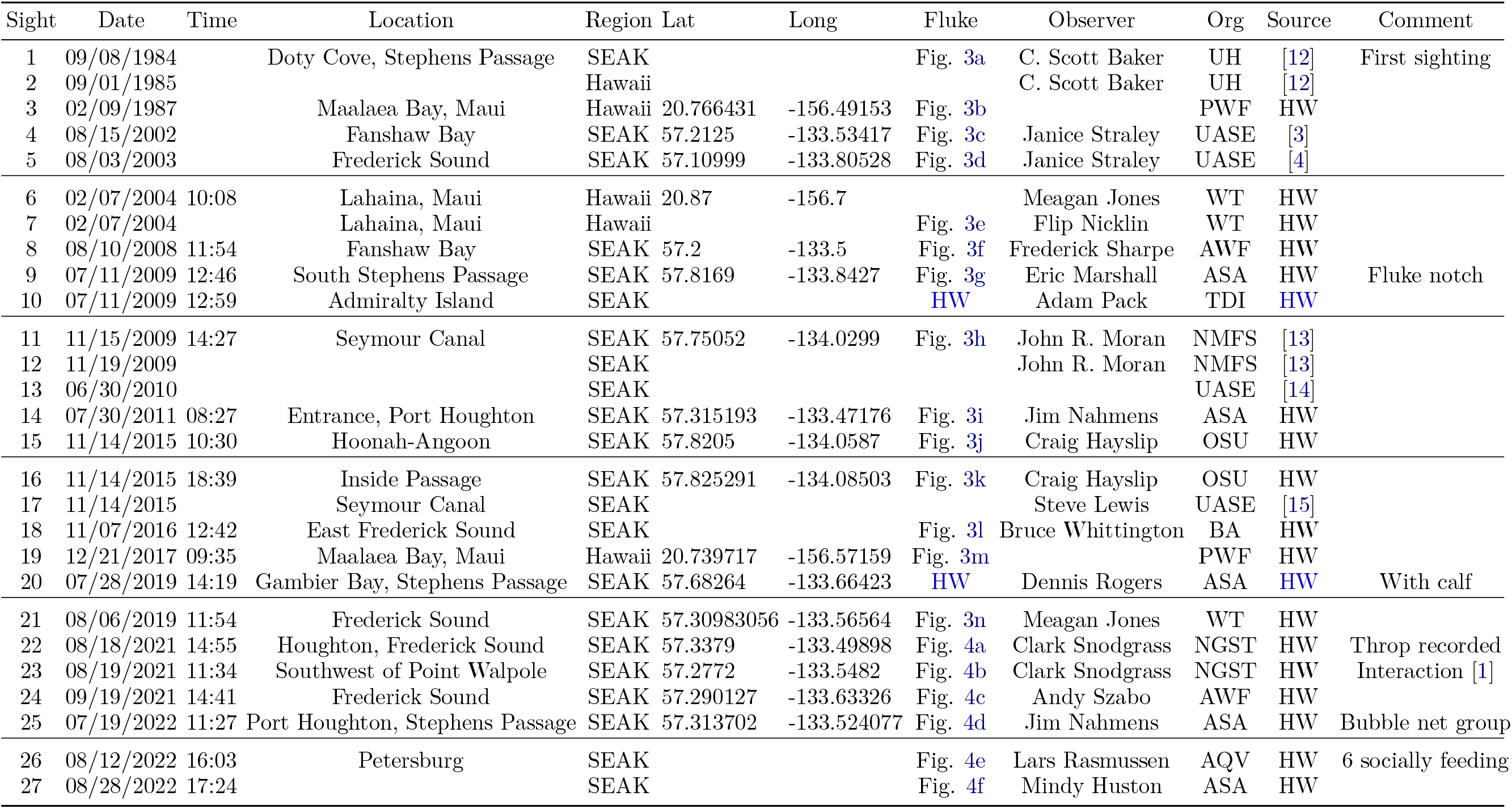
Chronology of Twain sightings.

**TABLE III:**
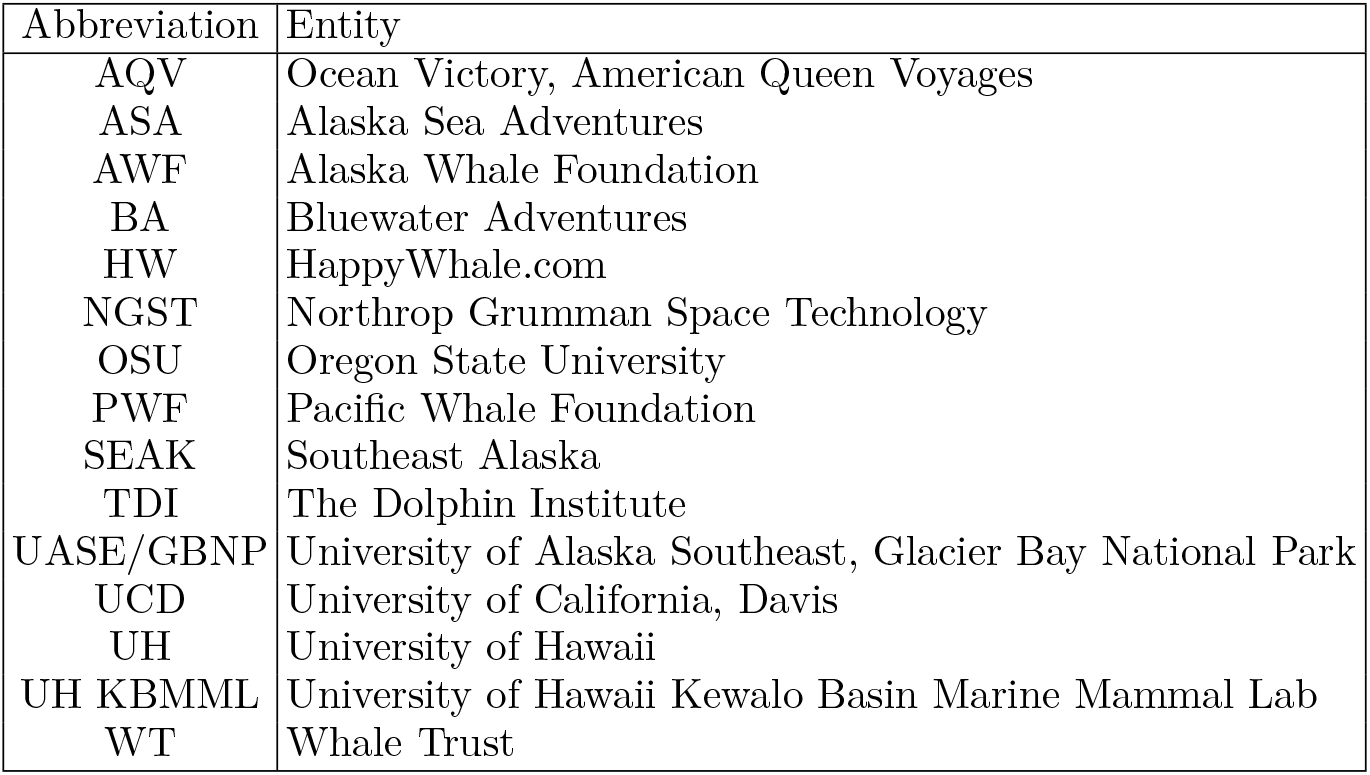
Abbreviations.

The majority of the sightings listed in Table II are available via HappyWhale.com and the UASE catalog. At the former, searching for “SEAK-0401” retrieves the majority of the sightings listed in Table II, including the documenting fluke photographs. Using the latter one can verify the animal’s identification for themselves, once a familiarity with the fluke markings is developed.

For the latter, in Twain’s case on the left of the fluke underside there are three diagonal linear-markings, with the middle marking consisting of three co-linear white circles. These allowed moderately rapid visual identification, even in the presence of highly-variable photographic quality and fluke orientation.

In addition to an ID label and fluke photograph, reported sightings typically, but not always, include date, time, and latitude and longitude of the encounter. Occasionally, there are circumstantial comments; see, for example Table II’s Comment column.

The earliest publicly-documented sighting, listed on HappyWhale, was in 1987; Fig. 3b. However, the first was by C. Scott Baker in 1984 in Southeast Alaska, Fig. 3a, and additional personal communications report a sighting by Baker a year later in 1985 in Hawaii [12]. Thus, the available documentation indicates that Twain is at least 38 years old and, certainly, even older. Given a speculated lifespan of 70 years the observation record covers the majority of Twain’s life.

The gallery presented here of almost four decades of fluke identifications does lead to further interesting observations and questions.

For example, the July 2019 sighting notes that Twain was seen with a calf. While the recent July and August 2022 sightings suggest that Twain was participating in bubble-net feeding groups.

In addition, the gallery also allows one to see how much Twain’s fluke markings changed. Encrustations— barnacles and vegetal growths—on her fluke tips clearly change, as expected. Also, between 2008 and 2009 Twain’s left fluke trailing edge took on a semi-circular divot. This is seen in the photos on the left fluke, close to and to the left of the fluke notch. This feature particularly helpful for identification in later years. It is clearly seen in all photos since 2009. See Fig. 5.

**FIG. 5:**
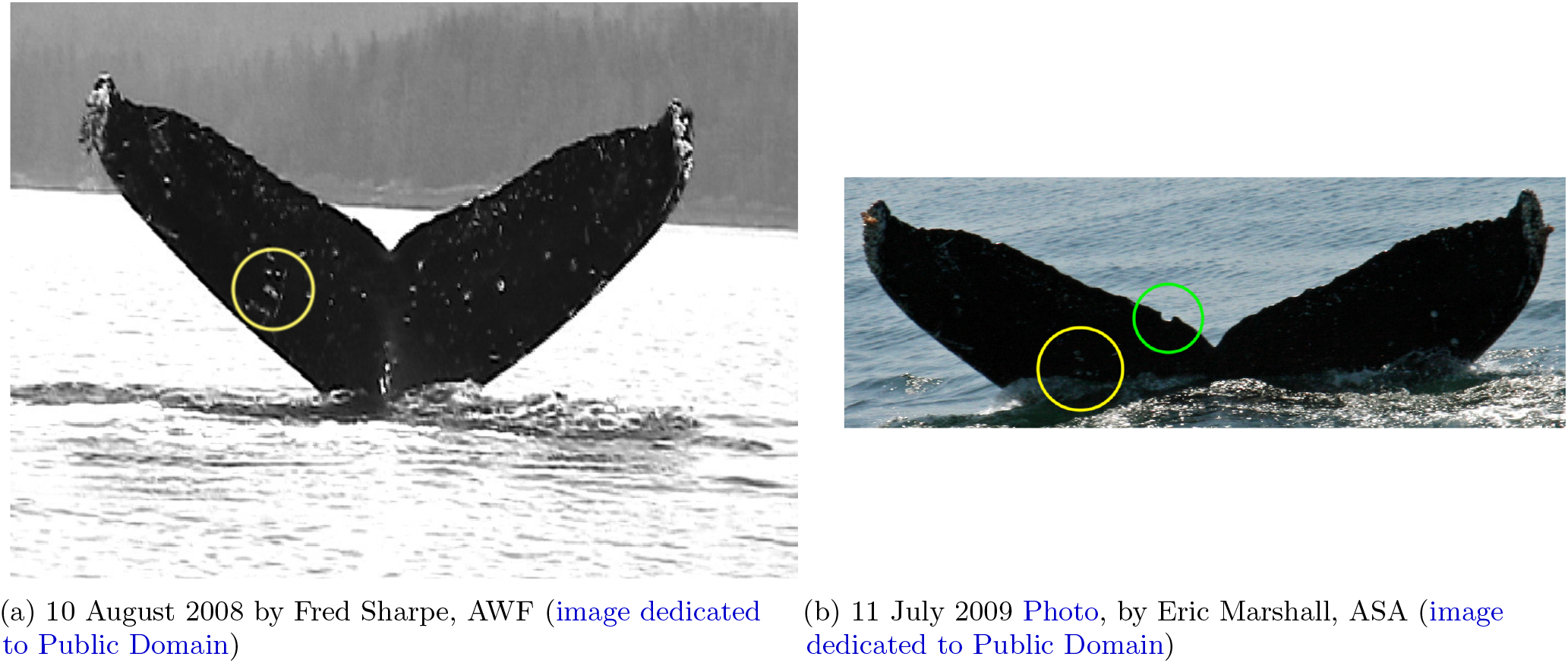
Evolution of Twain’s identifying fluke marks: (Left) In 2008 the characteristic three-parallel-line marks (yellow circle) are apparent. (Right) In 2009 these marks are seen again (dimly), but in addition a semi-circular divot appears on the left trailing fluke edge (green circle). Also, note the characteristic, for Twain, barnacle growth on the fluke tips.

## IV. CONCLUSION

Twain’s documented long life-history adds a new dimension to the recent extended acoustic exchanges reported in Ref. [1]. Rather than the latter being a singular encounter—and one unlikely to be repeated given the exigencies of very remote field work—it is now associated with an individual animal—a female, mother, widely traveled, with a record of health and interactions indelibly recorded on her fluke.

While there is still much to extract from the documented history and the acoustic interactions, a number of lessons have been learned and hopes for the future kindled.

The clear benefit of humpback fluke databases to appreciating a bit of Twain’s long life is unassailable. This leads, especially coming in the current setting motivated by acoustic interactions with Twain, one to advocate for expanded databases that include, for example, acoustic recordings of individual vocalizations, especially of identified individuals. This is certainly going to be necessary to make headway on understanding humpback social and song communication.

As desirable as these augmented databases will be, creating them presents a series of daunting tasks: from the shear serendipity required in the field to re-engage particular individuals—especially those like Twain who appear to have some motivation or interest in acoustically interacting, to the signal processing methods that will be required for automatic individual identification. These challenges seem to call for a radically new and greatly expanded research effort in cetacean biology and animal communication coordinated with mathematical, technical, and field innovations.

## ADDITIONAL INFORMATION

Correspondence and requests for materials should be addressed to the first author. The authors would very much appreciate new and also corrected information to help complete as much as possible of Twain’s life history.

## ACKNOWLEDGMENTS

Particular thanks to Scott Baker, John Moran, Fred Sharpe, Clark Snodgrass, and Andy Szabo for sharing observational data. J.P.C. thanks Brenda McCowan, Fred Sharpe, and Clark Snodgrass for helpful discussions. Some data was collected under NOAA permits issued to J. M. Straley. The prior field investigations noted were conducted under National Marine Fisheries Service Research Permit #19703 and partially funded by Templeton World Charity Foundation Diverse Intelligences grant TWCF0570 to University of California, Davis (Lead P.I. J. P. Crutchfield) and Templeton World Charity Foundation grant TWCF0440 to the SETI Institute (Lead P.I. L. Doyle; Co-PIs J. P. Crutchfield, M. Fournet, B. McCowan, and F. Sharpe).

## Author contributions

T.C., J.P.C., A.M.J., and J.M.S. collected the sighting data from the cited sources. J.P.C. and A.M.J. wrote and edited the manuscript and created the graphics.

## Data and materials availability

Data needed to evaluate the conclusions are available in the cited sources.

